# Age differences in brain white matter microstructure in UK Biobank (N = 3,513)

**DOI:** 10.1101/051771

**Authors:** Simon R. Cox, Stuart J. Ritchie, Elliot M. Tucker-Drob, David C. Liewald, Saskia P. Hagenaars, Gail Davies, Joanna M. Wardlaw, Catharine R Gale, Mark E. Bastin, Ian J. Deary

## Abstract

Quantifying the microstructural properties of the human brain’s connections is necessary for understanding normal ageing and disease states. We examined brain white matter MRI data in 3,513 generally healthy people aged 45-75 years from the UK Biobank sample. Using conventional water diffusion measures and newer, as-yet rarely-studied indices from neurite orientation dispersion and density imaging (NODDI), we document large age differences in white matter microstructure. Mean diffusivity was the most age-sensitive diffusion measure, with negative age associations strongest in the thalamic radiation and association fibres. Inter-individual differences in white matter microstructure across brain tracts become increasingly correlated in older age. This connectivity ‘de-differentiation’ may reflect an age-related aggregation of systemic detrimental effects on the brain. We report several other novel results, including comparative age associations with volumetric indices and associations with hemisphere and sex. Results from this unusually large, single-scanner sample provide one of the most definitive characterisations to date of age differences in major white matter tracts in the human brain.

**Abbreviations:**

**Tracts**
AR: acoustic radiation
ATR: anterior thalamic radiation
CingG: cingulum bundle; gyrus
CingPH: cingulum gyrus; parahippocampal
CST: corticospinal tract
FMaj: forceps major
FMin: forceps minor
IFOF: inferior fronto-occipital fasciculus
ILF: inferior longitudinal fasciculus
MCP: middle cerebellar peduncle
ML: medial lemniscus
PTR: posterior thalamic radiation
SLF: superior longitudinal fasciculus
STR: superior thalamic radiation
Unc: uncinate fasciculus

**Measures of white matter microstructure**
FA: fractional anisotropy
ICVF: intracellular volume fraction
ISOVF: isotropic volume fraction
MD: mean diffusivity
NODDI: neurite orientation dispersion and density imaging
OD: orientation dispersion

## Introduction

Fully understanding brain ageing requires an accurate characterisation of how and where white matter microstructure varies with age. White matter is highly relevant to ageing: later-life cognitive decline may partly be caused by ‘cortical disconnection’, a microstructural deterioration of the brain’s connective pathways through processes such as axonal demyelination that reduces information transfer efficiency^1–3^. The concept of disconnection has in large part been supported by analyses of diffusion MRI (dMRI); a non-invasive, quantitative method that exploits the Brownian motion of water molecules, allowing inferences to be made about the underlying microstructure of brain white matter *in vivo*^4–6^.

Past findings have been inconsistent in their characterisation of the trajectory and spatial distribution of age effects across brain white matter tracts, and of associations with hemisphere and biological sex^2, 7–23^. A statistically well-powered study of age differences in white matter microstructure, particularly in middle-aged and older age groups (45+ years), would address these gaps in our understanding. In addition to conventional measures of fractional anisotropy (FA; the directional coherence of water molecule diffusion) and mean diffusivity (MD; the magnitude of water molecule diffusion), newer and rarely-studied neurite orientation dispersion and density imaging (NODDI)^24^ measures offer new information on the microstructural bases of age effects on white matter^25^. NODDI provides estimates of neurite density (intra-cellular volume fraction; ICVF), extra-cellular water diffusion (isotropic volume fraction; ISOVF) and tract complexity/fanning (orientation dispersion; OD). The observed mean decline of FA in older age could be affected by, among other factors^5^, decreases in the density and/or an increase in the dispersion orientation of neurites (dendrites and axons); FA is unable to differentiate between these possibilities^26^. Thus, NODDI may offer a novel mechanistic insight into white matter ageing. However, a comprehensive examination of NODDI parameters has not been undertaken in the context of older age, and has not been attempted alongside more commonly-used water diffusion parameters.

The diffusion properties of white matter tracts across the brain are correlated; for example, an individual with relatively high FA in one tract is likely to also have relatively high FA across other tracts in the brain. This means that a latent, general factor of white matter microstructure can be derived^16, 27^. Analysis of latent factors allows a deeper understanding of the covariance structure of inter-individual differences in age-related brain changes; measuring global white matter diffusion is of great interest for investigating ageing trends that are general across white matter tracts (but specific to white matter tissue)^10, 11, 14, 21, 28^. Isolated, tract-specific enquiry cannot distinguish whether ageing patterns are unique to that tract, or an outcome of a more systemic constellation of processes. Tract-specific enquiry is also susceptible to a large degree of noise in the diffusion signal^29^; this noise can be reduced by using multivariate, latent-variable analyses^13^.

An additional gap in our understanding relates to the *de-differentiation* hypothesis, the suggestion that interindividual differences in microstructure across tracts become increasingly related in older age age^30–32^. This hypothesis stems from a “common cause” theory of ageing-related neurodegeneration^33^. It posits that system-wide breakdown of physiological function shifts overall levels of integrity across brain regions for affected individuals. Given that these sources of variance are shared across brain tracts, the signature should be higher correlations among regionally distributed neuronal measures with age. Here, we seek evidence for white matter structural de-differentiation from middle to older age, which could be an important marker of an aggregation of deleterious effects operating on white matter connections distributed across the central nervous system.

We undertook an analysis of age differences in 27 major white matter tracts (Figure 1). The data were from the UK Biobank resource (http://www.ukbiobank.ac.uk). We examined the tract-specific differences according to age, sex and hemisphere. We conducted analyses across all five water diffusion measures discussed above (FA, MD, ICVF, ISOVF, and OD), characterising which biomarker and which tract was most age-sensitive. We investigated whether tracts’ diffusion characteristics were more strongly correlated in older individuals (testing brain microstructural ‘de-differentiation’).

**Figure 1.**
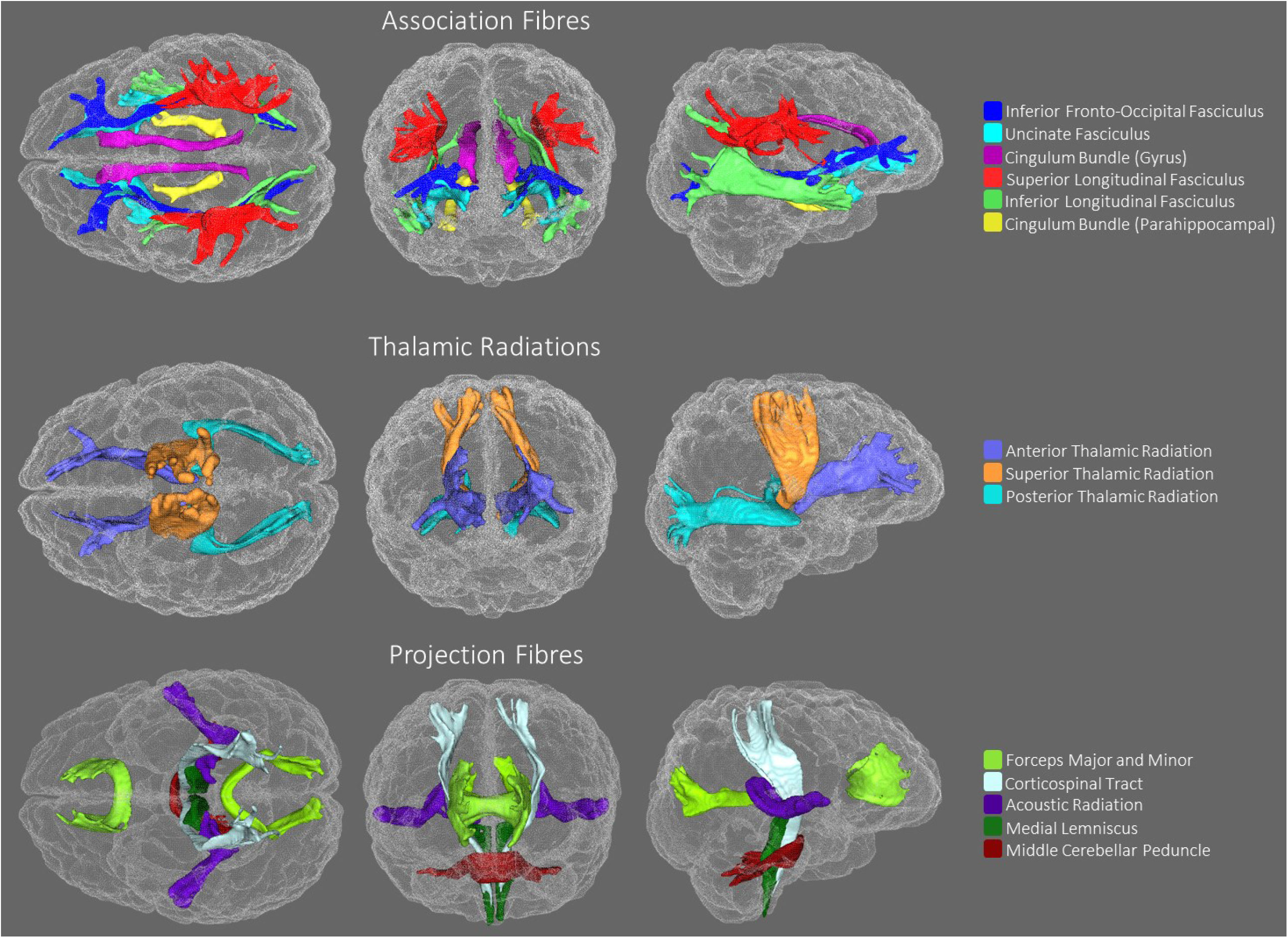
Tracts-of-interest, generated using probabilistic tractography rendered in superior (left), anterior (centre) and lateral (right) views.

## Results

### Tract-specific characteristics, and associations with age, hemisphere and sex

Characteristics of the 3,513 participants and a recruitment flowchart are reported in Supplementary Table 1 and Supplementary Figure 1. The current sample—the first release of the UK Biobank MRI sample—is a group of generally healthy middle-aged and older adults (age range 44.64 to 77.12 years). Tract-averaged values for MR diffusion parameters in each brain pathway are displayed in Supplementary Figure 2 and Supplementary Tables 1 and 2. Associations among left and right FA, MD, ICVF, ISOVF and OD values are displayed in Supplementary Figure 3 and 4.

Older age was significantly associated with lower coherence of water diffusion (FA; *β* ≥ −0.275), lower neurite density (ICVF; *β* ≥ −0.382) and lower tract complexity (OD; *β* ≥ −0.277), and with a higher magnitude of water diffusion (MD; *β* ≤ 0.496) and isotropic volume fraction (ISOVF; *β* ≤ 0.343) across the majority of tracts. These results indicate less healthy white matter microstructure with older age. Associations were predominantly non-linear with the exception of FA (Figures 2a–c; Supplementary Tables 3 and 4), indicating steepening slopes with increasing age. Associations with age were particularly marked in association (IFOF, ILF, SLF, Uncinate) and thalamic radiation (ATR, STR, PTR) fibres, as well as in the forceps minor (Figure 3). In contrast, the cingulum and sensory projection fibres showed modest or absent age differences across all diffusion parameters. Among these specific association and thalamic radiation fibres, MD exhibited the strongest associations with age of any of the diffusion parameters (Williams’s *t*-values > 4.27, *p*-values < 0.001; Figure 4).

**Figure 2a.**
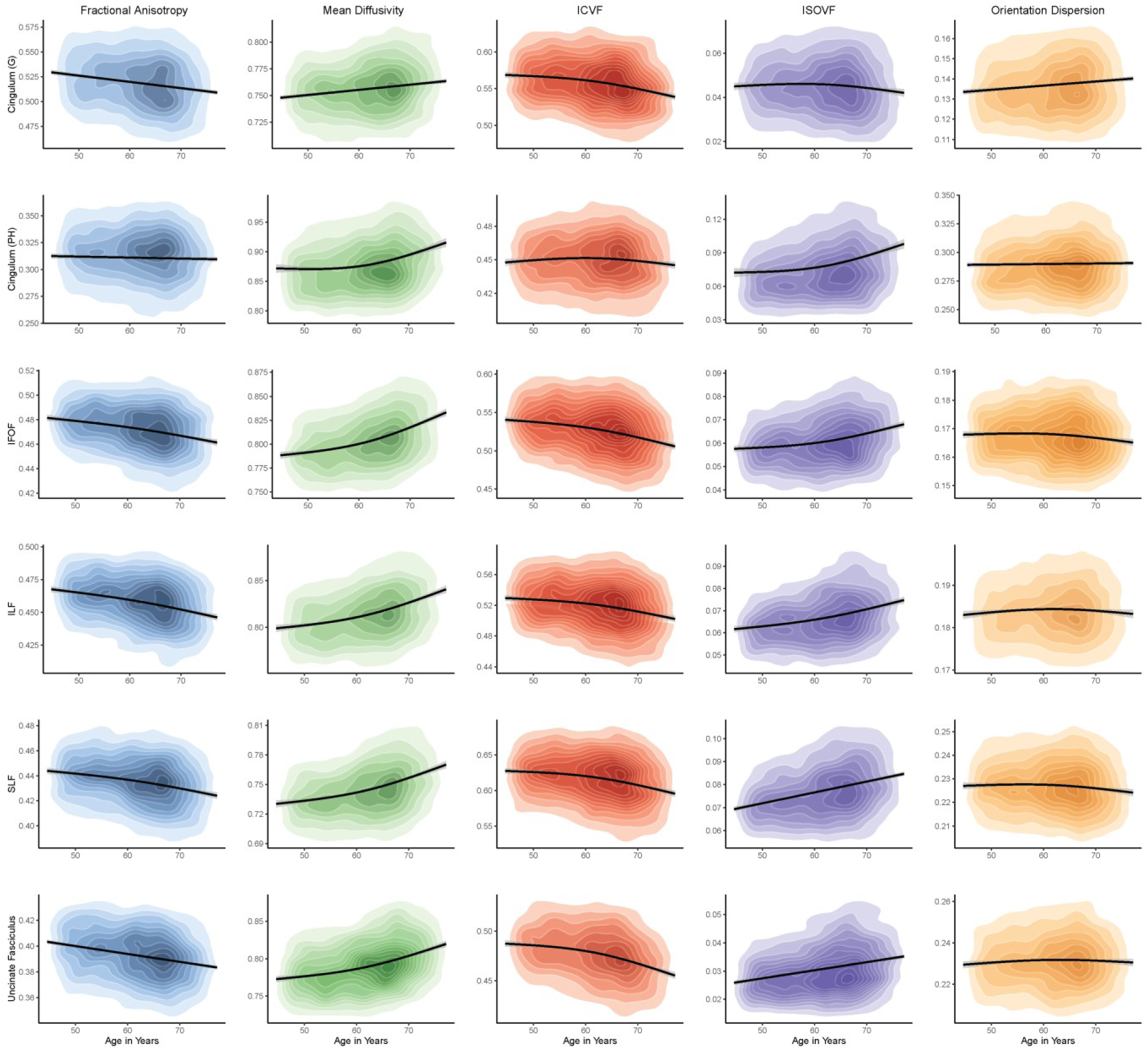
Age differences in the microstructural characteristics (y-axis) of association fibres. Kernel density plots indicate the degree of data point overlap (darker = greater); black line denotes the linear or quadratic regression line (with grey 95% CIs) across the five microstructural measures. Blue: fractional anisotropy, green: mean diffusivity, red: intracellular volume fraction, purple: isotropic volume fraction, orange: orientation dispersion. G: gyrus, PH: parahippocampal, IFOF: inferior fronto-occipital fasciculus, ILF: inferior longitudinal fasciculus, SLF: superior longitudinal fasciculus.

**Figure 2b.**
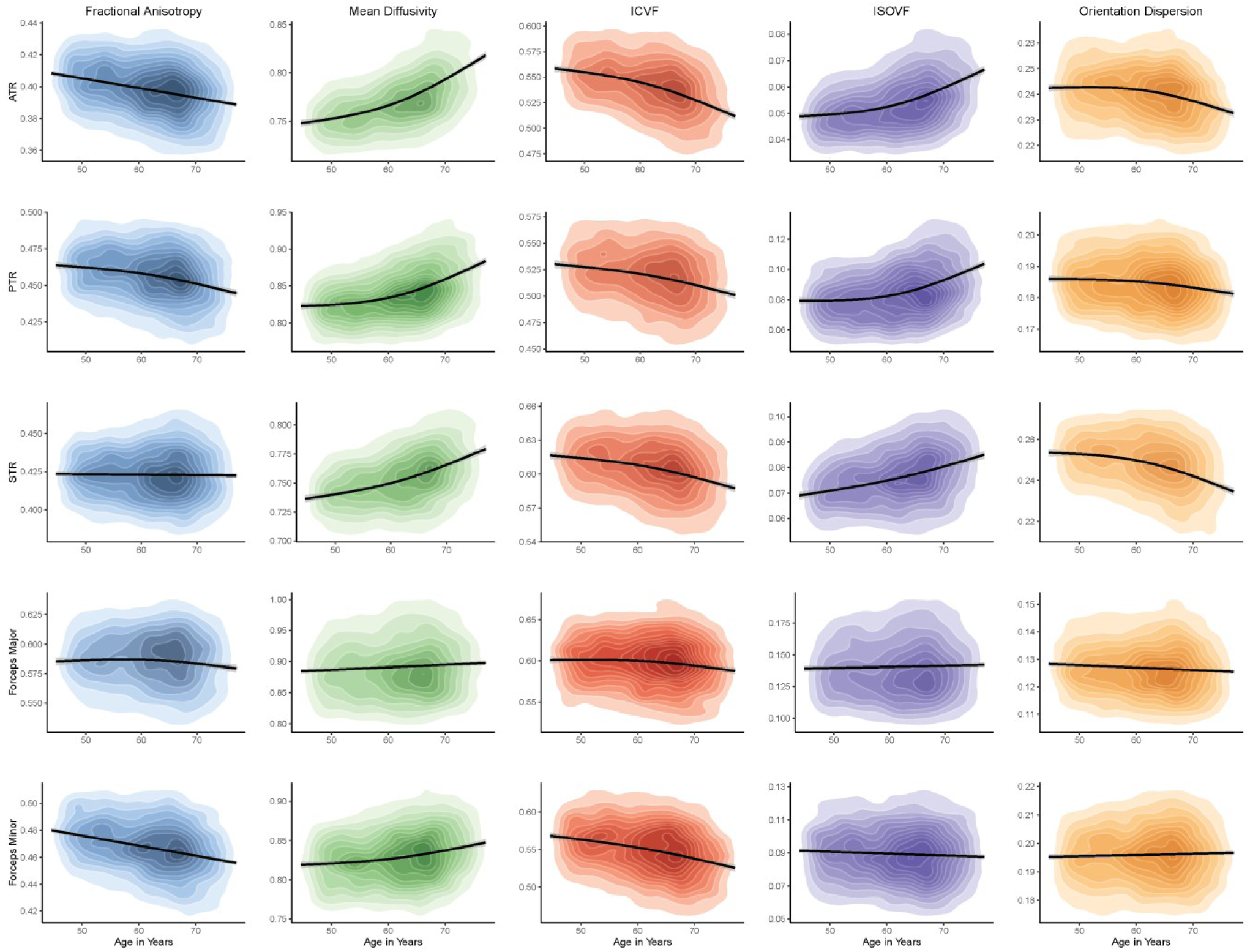
Age differences in the microstructural characteristics (y-axis) of thalamic and callosal fibres. Kernel density plots indicate the degree of data point overlap (darker = greater); black line denotes the linear or quadratic regression line (with grey 95% CIs) across the five microstructural measures. Blue: fractional anisotropy, green: mean diffusivity, red: intracellular volume fraction, purple: isotropic volume fraction, orange: orientation dispersion. ATR: anterior thalamic radiation, PTR: posterior thalamic radiation, STR: superior thalamic radiation.

**Figure 2c.**
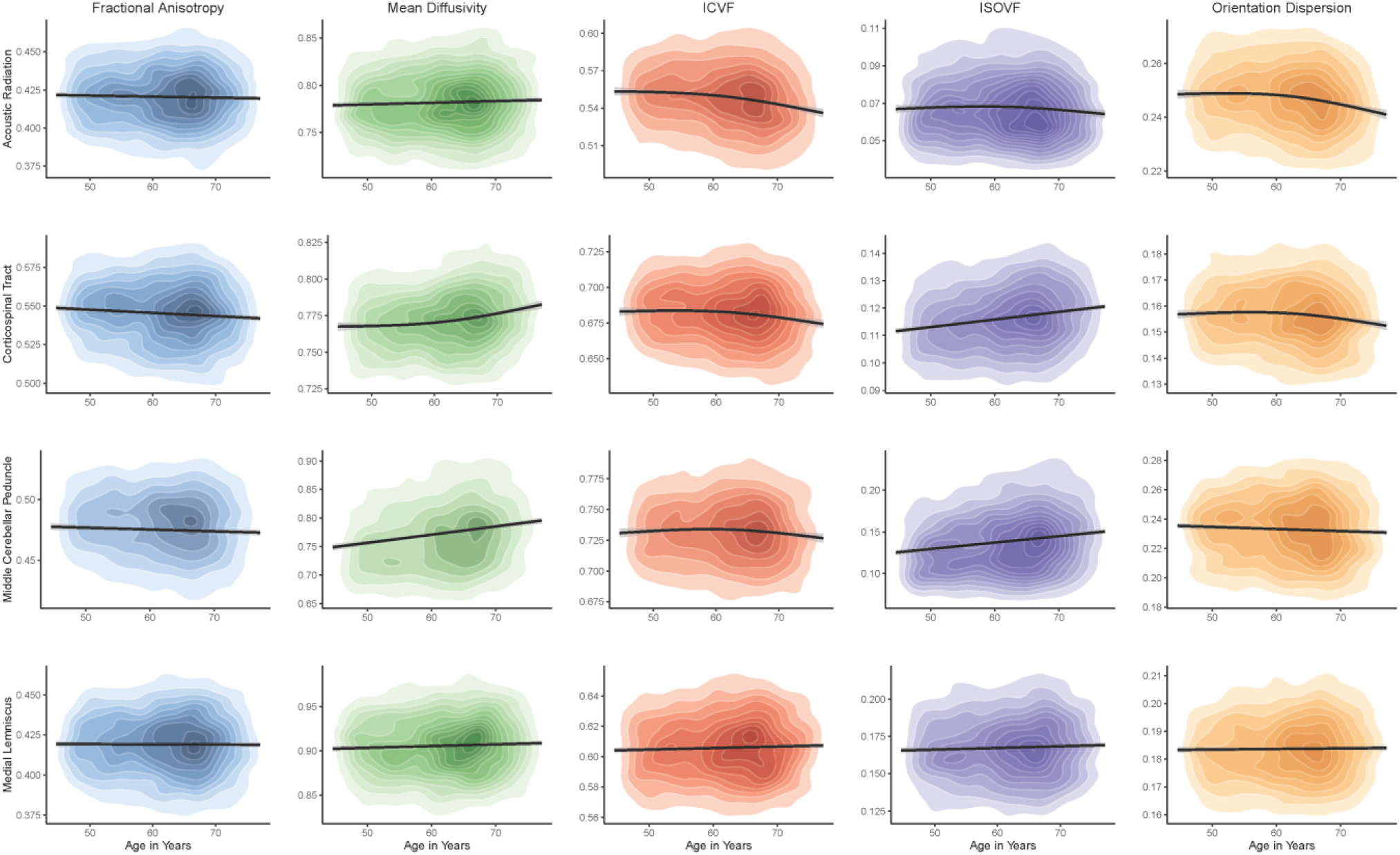
Age differences in the microstructural characteristics (y-axis) of sensory projection fibres. Kernel density plots indicate the degree of data point overlap (darker = greater); black line denotes the linear or quadratic regression line (with grey 95% CIs) across the five microstructural measures. Blue: fractional anisotropy, green: mean diffusivity, red: intracellular volume fraction, purple: isotropic volume fraction, orange: orientation dispersion.

**Figure 3.**
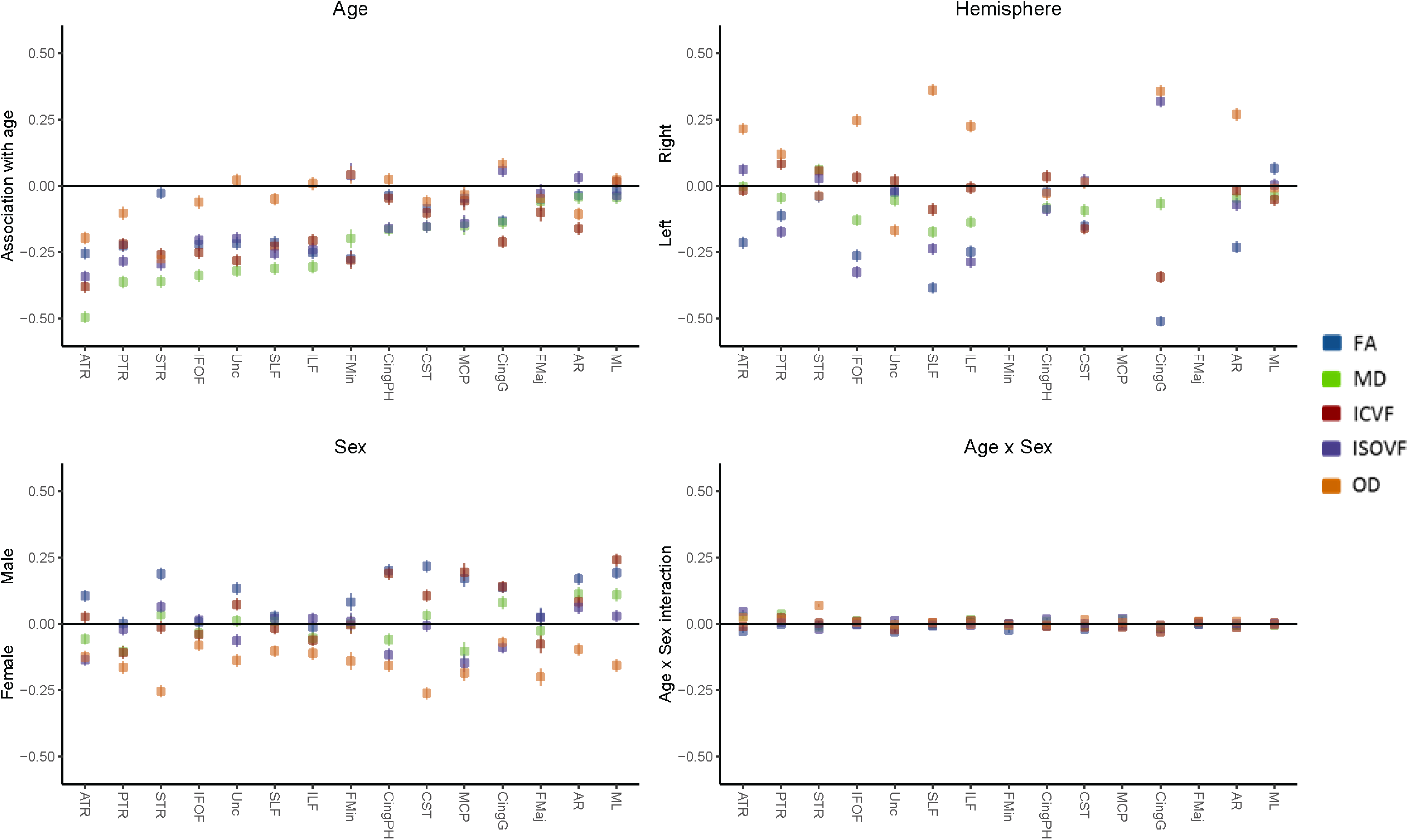
Results of regression analyses of tract-averaged FA (blue) MD (green), ICVF (red), ISOVF (purple) and OD (orange) with age, sex and hemisphere. Error bars = 95% CIs. Female and left hemisphere coded as 0. The valence of MD and ISOVF associations have been reflected for the purposes of visualisation for all four panels. See Supplementary Tables 3 and 4 for coefficients.

**Figure 4.**
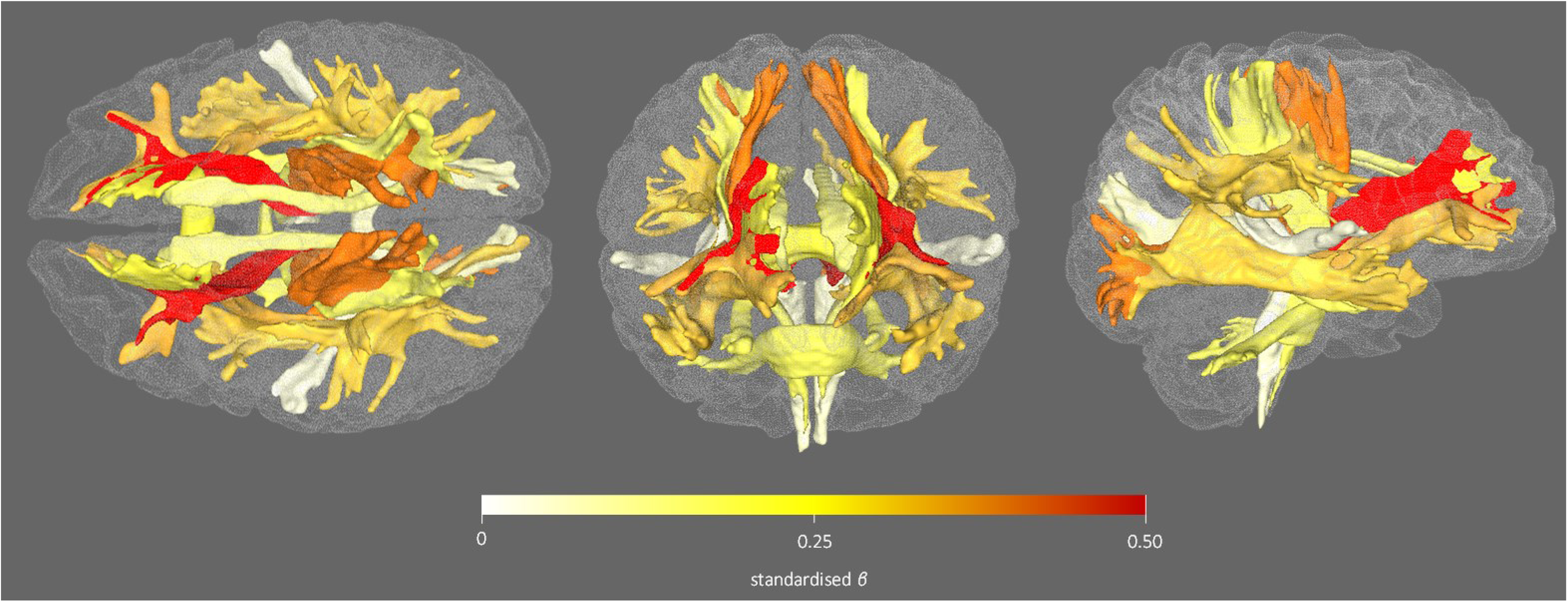
Illustration of association magnitudes (standardised *βs*) between age and tract-averaged MD. Rendered in superior (left), anterior (centre) and lateral (right) views. Coefficient values are from linear components of models shown in Supplementary Tables 3 and 4.

In addition to differential tract associations with age, putatively ‘healthier’ microstructure was found for some measures in the left versus right hemisphere, such as higher FA (*β* ≥ −0.511), lower ISOVF (*β* ≤ 0.319) and lower MD (*β* ≤ 0.175; Supplementary Tables 3 and 4, and Figure 3). Males also showed consistently higher FA (*β* ≤ 0.218). Females exhibited consistently greater OD (*β* ≥ −0.255). The interactions between age and sex—though significant for some tracts (age×sex *β*_*absolute*_ range = 0.028-0.079)—were small and inconsistent.

### General factors of white matter microstructure

We tested whether there was evidence for latent factors explaining a substantial portion of the variance in each of the five different types of tract measurement. That is, we tested whether a given microstructural measure was positively correlated among all tracts across the brain, and whether this common variance could be indexed by a latent factor variable.

Within all five white matter biomarkers, the measurements from across all tracts correlated positively; for instance, those with higher FA in one tract tended to have higher FA in all their tracts (Supplementary Figures 3 and 4). The ML, MCP, and the left/right CingPH tracts consistently covaried relatively weakly with others; these were removed from further models. The latent factors were thus indicated by 22 tracts each. For FA, MD, ICVF, and ISOVF, initial scree plots of the tract data (Figure 5, left panel) provided evidence for a strong single factor capturing common variance across the tracts; this was less clear for OD, which had a comparatively weaker first factor and a stronger second factor than the other measures. Across the age range, the first factor accounted for a mean of 41.4% of total variance in FA, 38.1% in MD, 68.2% in ICVF, 30.8% in ISOVF, and 20.1% in OD. Results presented below are therefore based on single factor models of each of the five diffusion measures. Henceforth, the prefix ‘*g*’ denotes these latent factors (e.g. the general fractional anisotropy factor is denoted ‘*g*FA’).

**Figure 5.**
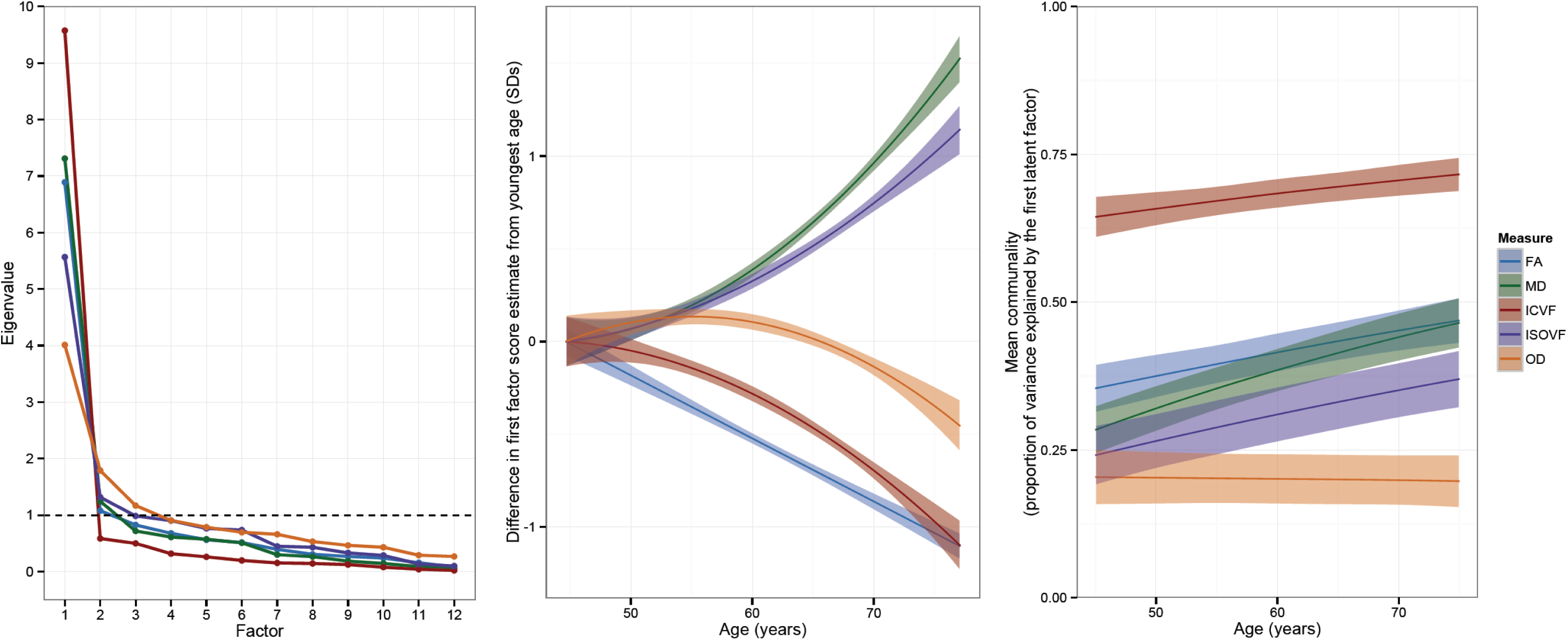
Left: Scree slopes for the exploratory factor analysis, showing the eigenvalue against the number of factors for each white matter tract measurement. Centre: Age trajectories of the first (latent) factor of white matter microstructure for each of the five dMRI biomarkers. Right: Age de-differentiation of white matter microstructure. Age trajectories for the proportion of total variance in each tract measurement explained by the general factor. The shaded region around each trajectory shows ±1 SD of the mean.

Age differences in the latent factors are illustrated in Figure 5 (central panel). As expected from the individual-tract data discussed above, *g*FA, *g*ICVF, and *g*OD factors were lower in older age. *g*FA and *g*ICVF showed relatively linear declines, and *g*OD showed decline after approximately age 60 years. *gMD* and *g*ISOVF showed substantial increases from 45 years of age. The standardised effect sizes for age were *g*FA: *β* = −0.25; *g*MD: *β* = 0.37; *g*ICVF: *β* = −0.27; *g*ISOVF: *β* = 0.27; *g*OD. *β* = −0.12 (Supplementary Table 7). Age had a significantly stronger association with *gMD* than with any of the other latent factors (all Williams’s *t*-values > 5.55, all *p*-values < 0.0001).

We next examined the extent to which the effect of age on white matter is common to all tracts, or tract-specific. We tested whether age associations with the individual tracts were accounted for by the association between age and the general factor (a ‘common pathway’), or whether there were incremental tract-specific age associations (‘common plus independent pathways’). For all five white matter microstructural measurements, there was evidence of age being associated with the general factor. There were also additional tract-specific effects; that is, ‘common plus independent pathways’ models fit significantly better than models that only included a ‘common pathway’ of age associations (Supplementary Table 8). In summary, age appears to affect the white matter both overall and in some additional, specific tracts (Supplementary Figure 5).

The FA signal is susceptible to microstructural properties in the brain that include neurite density and tract complexity/fanning. The availability of the NODDI variables ICVF and OD allow us to directly test if and how ICVF and OD mediate the association between increasing age and lower FA. We used estimates of the general factor scores in a multiple-mediator analysis. Our results (Supplementary Table 9 and Supplementary Figure 6) showed that the linear association between age and *gFA* was significantly mediated by 75.71% (from *β* = −0.247 to *β* = −0.060) with the inclusion of the NODDI parameters. The majority of this mediation took place through *g*ICVF rather than *g*OD. This suggests that age-related declines in brain white matter FA may predominantly be explained by declines in neurite density rather than changes in tract complexity.

Next, we aimed to contextualise the utility of all five white matter tract diffusion parameters’ general factors in a reverse inference exercise, examining how much age variation they could explain beyond conventional brain volumetric measures in the same participants (total brain, grey matter, white matter and bilateral hippocampal volumes; corrected for head size). Increasing age was associated with lower volumes in each of these measures (Supplementary Table 10 and Supplementary Figure 7). Given the high collinearity of all volumetric and diffusion indices (Supplementary Table 11), we employed a penalised (elastic net) regression to identify an optimal set of predictors of chronological age in one half of the randomly and equally split sample (training set).

The results showed that age-related variance in specific aspects of white matter microstructure (FA, MD and ISOVF) is partially independent of atrophy and grey matter volume. Of the five white matter microstructural measures and five volumetric measures, *g*FA, *g*MD, *g*ISOVF, total brain volume and grey matter volume appeared in more than 60% of the 1000 bootstrapped models on the training set (the remaining variables did not appear once). These predictors were entered into a multiple linear regression in the training set, followed by a confirmatory test using the same predictors in the testing set (Supplementary Table 12). In both cases, *g*FA (*β*_*train*_ = −0.085 and *β*_*test*_ = −0.071, *p* ≤ 0.035), *g*MD (*β*_*train*_ = 0.085 and *β*_*test*_= 0.109, *p* ≤ 0.019) and *g*ISOVF (*β*_*train*_ = 0.119 and *β*_*test*_ = 0.124, *p* < 0.001) accounted for unique age variance, beyond total brain volume (*β*_*train*_ = −0.160 and *β*_*test*_ = −0.171, *p* < 0.001) and grey matter volume (*β*_*train*_ = −0.375 and *β*_*test*_ = −0.348, *p* < 0.001). For both models, these predictors accounted for nearly 40% of the age variance (*R*^*2*^ = 0.377 in both cases), and did not exhibit multicollinearity (variance inflation factors < 3.84), indicating that the elastic net method had been effective in producing a useful set of predictor variables.

### Age de-differentiation of white matter microstructure

Do white matter tracts tend to lose their individuality in older age? We tested the ‘de-differentiation’ hypothesis, that white matter tracts across the brain become are more similar in their quality at later ages. We extended each of the five general factor models to include an interaction parameter representing age moderation of the shared variance across the 22 tracts and an interaction parameter representing age moderation of the tract-specific unique variance. In other words, we estimated two interaction parameters for each of the 5 diffusion measures. This allowed us, for each diffusion measure, to calculate the mean communality (the proportion of total variance that was shared) across tracts as a continuous function of age.

For four of the five diffusion measures, the latent factor accounted for more variance in older age (Figure 5, right panel). For FA, the factor explained 11.5% more variance in the oldest participants (around age 75) than in the youngest participants (around age 45). The equivalent differences were 18.1% for MD, 7.2% for ICVF, and 12.9% for ISOVF. There was no appreciable age difference in the first factor of OD with age (there was a small, non-significant decline in explained variance of −0.7%); we saw above that OD also evinced the weakest general factor overall. In summary, for four of the biomarkers, there was greater generality across the tracts with greater age, providing clear evidence of age-related de-differentiation. The older the brain, the more likely it will be that—within a given individual—one tract will share a microstructural quality with of all of the brain’s other white matter tracts.

Finally, we evaluated whether the overall pattern of increasing tract communality with age was driven by particular subsets of the tracts. These analyses (Supplementary Tables 13 and 14; Supplementary Figures 8-12) revealed that much of the upward mean trend in tract generality (for all measures except OD) shown in Figure 5 was driven by steeper trends in the communalities of the association fibres and thalamic radiations – the same tracts that showed the largest mean age differences. The communalities of the sensory projections (AR and CST) were flatter, indicating that the variance in them explained by the general factor was similar across the age range. Notably, some tracts (such as the Forceps Major) showed age differences in the amounts of variance explained at both the general and the specific level: that is, the total amount of tract variance was higher in older age.

Overall, we found that, at older ages, there was a greater tendency for all tracts in the brain to show similar levels of FA, MD, ICVF and ISOVF within individuals. The increases in the tracts’ common variance were particularly pronounced for the thalamic radiations and association fibres. Therefore, it was not simply that age differences were largest among these specific tracts, but that interindividual differences in diffusion measures for these tracts covaried more strongly with each other and with other tracts in the brain in older individuals.

## Discussion

This study adds to our understanding of the white matter microstructure of the brain in middle-and older-aged humans. Older age was most strongly associated with less healthy white matter in the thalamic radiations (ATR, PTR and STR) and association fasciculi (ILF, IFOF, SLF and Uncinate). The comparison of conventional dMRI and NODDI variables showed that MD was the most sensitive parameter to age among these tracts. Interindividual differences in each diffusion measure showed a clear tendency to covary across white matter tracts. General factors taken across the tracts captured substantial variation in FA, MD, ICVF and ISOVF, and more modest proportions of variation in OD. The general factors each had a tendency toward less healthy values at older ages, and we also observed tract-specific age associations beyond those that could be accounted for exclusively via general factors.

Mediation models indicated that the link between older age and lower general fractional anisotropy (*g*FA) was predominantly driven by lower neurite density (*g*ICVF) rather than greater orientation dispersion (*g*OD). In a reverse inference exercise, we illustrated the importance of brain diffusion parameters for understanding ageing: latent measures of white matter diffusion (FA, MD and ISOVF, in particular) provided unique information about age, beyond conventional volumetric brain measures of global atrophy, hippocampal, grey and white matter volume. Finally, the tendency for white matter tracts within individuals to share microstructural qualities (assessed using water diffusion characteristics) was stronger in older participants, indicating de-differentiation of brain connectivity. This tendency was most strongly driven by the greater relatedness among the same thalamic radiation and association fasciculi that showed the greatest age differences in our tract-specific analyses.

The apparent differential age trends—both heterochronicity and spatial heterogeneity—across the tracts is supportive of the ‘last in, first out’ hypothesis^34^, whereby the tracts that are latest to develop are the most vulnerable to the deleterious effects of ageing. In particular, post-mortem and dMRI studies indicate ontogenetic differences between early-myelinating projection and posterior callosal fibres, and the later-developing association pathways^12, 21, 22, 35^. An underlying reason for this pattern may be that the pathways latest to develop are more thinly myelinated, and that the oligodendrocites responsible for their myelination are more vulnerable to aggregating deleterious effects due to their comparatively elevated metabolic activity^2, 17^. We found that this pattern also occurs for more rarely-studied water molecule diffusion measures (ICVF and ISOVF). Thus, these spatially distinct relations with age are apparent across a wider range of microstructural properties than was previously thought.

Our finding that the strongest age associations were with the thalamic radiations does not, however, follow the reported ontogeny in childhood. Whereas the anterior limb of the internal capsule (which contains, among other fibres, the anterior thalamic radiation) shows a relatively delayed maturational trajectory, the posterior limb of the internal capsule (which contains, among other fibres, the posterior thalamic radiation) shows comparatively early development along with the cortisospinal tract, acoustic radiations and middle cerebellar peduncle^36, 37^, yet the PTR showed some of the most marked age differences in this sample. However, the initial supposition of ‘last in, first out’ referred to both ontogenetic and phylogenetic chronology^34^; the thalamic nuclei are likely to have undergone considerable evolutionary modification, perhaps to keep pace with the rapidly expanding cortex^38, 39^. The complex set of nuclei that comprise the thalamus share connections across the whole cortex, including hippocampal and prefrontal pathways, forming a densely interconnected processing unit^40, 42^. The current study highlights the potential importance of the anterior, superior and posterior thalamic radiations and association fibres for understanding brain ageing; these tracts could be foci of future investigations into behavioural outcomes and possible determinants.

Age-related differences in white matter diffusion measures were not simply reflective of gross volumetric brain indices in our sample. All diffusion and volumetric measures were, in isolation, associated with age, but our analysis identified that grey matter volume, total brain volume, and *g*ISOVF, *g*FA and *g*MD were significant predictors (together explaining ˜40% of the variance in age), whereas white matter volume, bilateral hippocampal volume, *g*ICVF, and *g*OD were not. The result that information about white matter water diffusion from both NODDI and more conventional measures is more informative for age than white matter volume might indicate that these biomarkers are more sensitive to subtle age-related differences in white matter. However, it is important to note that a global white matter measure can be divided into normal-appearing, and white-matter hyperintensity (WMH) volume in older subjects^43, 44^. The latter, a marker of white matter disease^43^, appears to progress independently of grey matter changes^45^, but was not derived in the current dataset. It may be that the insensitivity of total white matter volume to the important ratio of normal-appearing: WMH volume may partly account for its lack of predictive value in our models. Additionally, the current investigation focussed on specific pathways, therefore excluding peripheral white matter, which may also prove to be of particular interest to brain and cognitive ageing^46^.

The participants in the current study were in relative good health; we excluded individuals with any known neurodegenerative disease. Whereas there is evidence that MD is also more sensitive to age than FA among those with mild stroke^47^, our sample composition has implications for another aspect of our results. Whereas hippocampal volume and ICVF did not appear to provide unique information about age in this sample, both ICVF (rather than FA)^48^ and hippocampal volume^49^ are important dementia biomarkers, emphasising that our findings, which focus on normal ageing, cannot necessarily be generalised to clinical populations.

From each of the five white matter parameters, we extracted latent general factors to be extracted from the covariances among 22 white matter tracts. These factors indexed from 20% to 68% of the variance across the tracts, highlighting their importance—but not exclusivity—for providing indices of overall brain microstructure. The fit of these models was improved by adding correlated residual paths beforehand, and we removed three fibre pathways for having low factor loadings. Thus, these are not the *only* factors that were extractable from the data: smaller, weaker factors may also exist that index appreciable portions of cross-tract variance^16^. Consequently, whereas we show that the association between age and *g*FA was predominantly attenuated by *g*ICVF rather than *g*OD, we note that the latter did not exhibit such a strong first factor, and therefore it is possible that our estimate of the amount that *g*OD contributes to the age-*g*FA relation might fluctuate more as a function of tract than it might for *g*ICVF.

In our analysis of ‘de-differentiation’ in the brain, we found compelling evidence—in four of the five white matter microstructural measurements (all except OD)—for increasing tract generality with age. For each of these four microstructural properties, the covariation across tracts within the brain was higher with greater age. One explanation of this de-differentiation is that the ageing of white matter structure involves an aggregation of systemic, detrimental effects that render tract-specific variation in white matter (dis)connectivity less prominent at older ages. Uncovering the specific cellular and metabolic mechanisms that might cause this increasing generality, and how this generality might shed light on the brain basis for cognitive and functional ageing, should be investigated in future studies.

Further to limitations of our sample composition and the brain MRI and statistical analyses discussed above, cross-sectional estimates of age-related trajectories may imperfectly index patterns of within-person changes over time^50^. They are also subject to bias due to cohort differences and secular trends in brain structure. Nevertheless, longitudinal studies remain rare and the time course required to test healthy participants prospectively across a comparable age span is prohibitive, leading many to adopt semi-longitudinal designs across large age ranges^8, 10, 16, 23, 51, 52^, with comparatively smaller samples across a wide age range, or adopt full longitudinal designs in cohorts with a narrow age range^3^. Thus, while the current cross-sectional data provide a well-powered insight into differences in white matter microstructure with age, examination of intra-individual change awaits further data. Fully longitudinal studies in larger, wide-age-span samples would be required to track individual trajectories directly, though coverage of a comparable time span to the current study would take many years.

This large-scale, single-scanner brain imaging sample has afforded clear insights into the human brain’s connections in middle to older age. In this paper we: located and quantified the age-related differences in white matter tracts; provided robust information about which diffusion-based biomarkers were especially sensitive in ageing; demonstrated that the inter-individual variation in specific tracts became less specific with age; and found that ageing processes are best modelled as acting upon those characteristics that tracts share rather than those that are unique to them. Thus, whereas we found that the brain is more ‘disconnected’ in later life^1^ (since white matter microstructure was in poorer health), in another sense we found it to be increasingly ‘connected’, since its tracts were more similar at older ages. These findings offer secure foundations for planned further exploration of the risk factors and mechanisms of brain and cognitive ageing.

## Online Methods

### Ethical Approval, Data Availability and Consent

Researchers can apply to use the UK Biobank data resource for health related research in the public interest. A guide to access is available from the UK Biobank website (http://www.ukbiobank.ac.uk/register-apply/). UK Biobank received ethical approval from the research ethics committee (REC reference 11/NW/0382). The present analyses were conducted under UK Biobank application number 10279. All participants provided informed consent to participate. Further information on the consent procedure can be found here (http://biobank.ctsu.ox.ac.uk/crystal/field.cgi?id=200).

### Participants

The UK Biobank comprises ˜500,000 community-dwelling participants who were initially recruited from across Great Britain between 2006 and 2010, aged 40-69 years (http://www.ukbiobank.ac.uk). An average of 4.15 (SD = 0.91) years after initial recruitment, a subset of participants also underwent head MRI at mean age 61.72 (SD = 7.47, range 44.64-77.12) years. The initial release of brain diffusion MRI (dMRI) data from 5,455 participants is the subject of the current study.

### Demographic Information

Information on qualifications, ethnicity, sex and handedness were reported during the initial UK Biobank assessment (http://biobank.ctsu.ox.ac.uk/crystal/refer.cgi?id=100235). Educational qualifications were taken from responses to the question: “Which of the following qualifications do you have? (You can select more than one)”. Response options were: College or University Degree / A levels or AS levels or equivalent / CSEs or equivalent / NVQ or HND or HNC or equivalent / Other professional qualifications e.g. nursing, teaching / None of the above / Prefer not to answer. For the purposes of characterising the participants here, we collapsed the data into a binary variable indicating whether or not each participant held a college or university degree. Self-reported ethnic background was based on response to the question “What is your ethnic group?”. Response options were: White / Mixed / Asian or Asian British / Black or Black British / Chinese / Other ethnic group / Do not know / Prefer not to answer. These responses were collapsed into White, Mixed and Other. Handedness was based on responses to the question: “Are you right or left handed”, where response options were: Right-handed / Left-handed / Use both right and left equally / Prefer not to answer. At the time of the MRI assessment, participants’ medical history was taken and coded by a trained nurse according to a specific coding tree (http://biobank.ctsu.ox.ac.uk/crystal/field.cgi?id=20002). Those who reported a diagnosis of dementia/alzheimer’s disease or mild cognitive impairment, Parkinson’s disease, stroke, other chronic/degenerative neurological problem or demyelinating condition (including multiple sclerosis and Guillain-Barré syndrome) were removed from analysis.

### MRI Acquisition

Details of the image acquisition are freely-available on the UK Biobank website in the form of a Protocol (http://biobank.ctsu.ox.ac.uk/crystal/refer.cgi?id=2367) and Brain Imaging Documentation (http://biobank.ctsu.ox.ac.uk/crystal/refer.cgi?id=1977). Briefly, all brain MRI data were acquired on a single standard Siemens Skyra 3T scanner with a standard Siemens 32-channel RF receiver head coil, with the imaging matrix angled down by 16° from the AC-PC line. The T1-weighted volumes were acquired in the sagittal plane using a 3D MPRAGE sequence at a resolution of 1 × 1 × 1 mm, with a 208 × 256 × 256 field of view (FoV). The dMRI protocol employed a spin-echo echo-planar imaging (SE-EPI) sequence with 10 T2-weighted (b ≈ 0 s mm^−2^) baseline, 50 b = 1000 s mm^−2^ and 50 b = 2000 s mm^−2^ diffusion-weighted volumes acquired with 100 distinct diffusion-encoding directions and three times multi-slice acquisition. The FoV was 104 × 104 mm, imaging matrix 52 × 52, 72 slices with slice thickness 2 mm, giving 2 mm isotropic voxels. The flowchart in Figure S1 illustrates the numbers from initial MRI recruitment and attendance through to completion and quality control procedures. Of the 5,455 who provided MRI data, 567 were acquired at an earlier scanning phase (for which the resultant dMRI data are incompatible with data acquired subsequently; see Section 2.10 of the Brain Imaging Documentation). A further 1,314 participants were removed during dMRI quality control procedures by UK Biobank prior to data release, which was a combination of manual and automated checking and also included the removal of data badly affected by movement artefacts (as described in the UK Biobank Brain Imaging Documentation). In addition to the 59 participants with self-reported diagnosis of stroke, dementia, Parkinson’s disease, or any other demyelinating or neurodegenerative disorder, a further 2 participants with consistently extreme outlying tract-averaged water diffusion biomarker values were removed listwise, along with 35 individual extreme outlying data points (< 0.001% of total data), following visual inspection of the data by the authors, leaving a total of 3,513 participants for analysis in the current study. The current sample did not differ from those not scanned with respect to age at initial recruitment (*t* = 0.834, *p* = 0.405), but comprised a higher proportion of females (*χ*^2^ = 9.637, p = .002). The total numbers of available tracts are reported in Supplementary Tables S1 and S2.

### Diffusion MRI Processing and Tractography

Fractional anisotropy (FA) and mean diffusivity (MD) are commonly-derived variables which describe the directional coherence and magnitude of water molecule diffusion, respectively. Water molecules tend to diffuse with greater directional coherence and lower magnitude when constrained by tightly-packed fibres (such as well-myelinated axons) as well as by cell membranes, microtubules and other structures^26^. Thus individual differences in FA and MD in brain white matter reflect meaningful differences in underlying microstructure, borne out by comparison to brain white matter post-mortem work^4, 5^. Unlike standard diffusion tensor MRI, NODDI makes specific assumptions about the way in which local microarchitecture affects the geometric diffusion of water, and parameterises the water diffusion signal according to one of three geometrical models: free water diffusion (such as in CSF), restricted diffusion caused by the presence of dendrite and axons bodies, and hindered diffusion among cell bodies. The resultant indices describe the intracellular volume fraction (ICVF; a measure of neurite density), isotropic volume fraction (ISOVF; a measure of extracellular water diffusion), and neurite orientation dispersion (OD; the degree of fanning or angular variation in neurite orientation).

Gradient distortion correction was applied using tools developed by the Freesurfer and Human Connectome Project groups, available at https://github.com/Washington-University/Pipelines. The Eddy tool from FSL (http://fsl.fmrib.ox.ac.uk/fsl/fsiwiki/EDDY) was then used to correct the data for head motion and eddy currents. Next, within-voxel multi-fibre tract orientation structure was modelled using BEDPOSTx followed by probabilistic tractography (with crossing fibre modelling) using PROBTRACKx^53–55^. Automatic mapping of 27 major tracts in standard space using start/stop ROI masks (implemented using the AutoPtx plugin for FSL)^56^ was then use to derive tract-averaged measures of FA and MD for the following tracts of interest: middle cerebellar peduncle, forceps major, forceps minor, and bilateral medial lemnisci, corticospinal tracts, acoustic, anterior thalamic, posterior thalamic, superior thalamic radiations, superior, inferior longitudinal and inferior fironto-occipital fasciculi, and both the cingulate gyrus and parahippocampal portions of the cingulum bundle (Figure 1 in the main document). In addition, NODDI (Neurite Orientation Dispersion and Density Imaging)^24^ modelling of the dMRI data was conducted using the AMICO tool (Accelerated Microstructure Imaging via Convex Optimization; https://github.com/daducci/AMICO)^57^. Maps of ICVF, ISOVF and OD, registered with the AutoPtx tract masks, allowed the calculation of tract-averaged values for each parameter across all voxels pertaining to each tract of interest.

### Volumetric MRI Processing

Extraction of the brain was achieved by non-linearly warping the data to MNI152 space (FNIRT)^58, 59^, with the brain mask then back-transformed into native space. FAST^60^ was then used to segment the brain tissue (in native space, to avoid noise due to interpolation) into CSF, grey matter and white matter (with total brain volume being the sum of grey and white matter volume); bilateral hippocampal volumes were derived using FIRST^61^. All volumes were then adjusted for head size using a SIENAX-style analysis^62^. This involves deriving a scaling factor from the normalisation transform matrix obtained from the affine registration of skull tissue between T1-weighted volume and MNI152 space. The resultant scaling factor was then applied to the volumes of interest for each participant.

### Statistical Analysis

Statistical analyses were conducted using R v3.2.2 (‘Fire Safety’) and MPlus v7.3^63^. The distribution of each white matter tract diffusion measure was inspected, and we found no large deviations from normality.

Handedness is sometimes reported in dMRI studies, under the assumption that differences exist (e.g. Bender et al.^8^). There were no substantive differences in tract characteristics between left-and right-handed participants (except for a trend for left-handers to have marginally higher FA (*t* (378.53) = 2.130, *p* = 0.034, *d* = 0.219) and lower OD (*t* (374.58) = −2.747, *p* = 0.006, *d* = 0.141) in the forceps minor, lower OD in the left CST (*t* (378.19) = −2.446, *p* = 0.015, *d* = 0.125) and higher OD in the right Uncinate (*t* (373.85) = 2.297, *p* = 0.022, *d* = 0.238). As a result, all analyses are reported across the entire group. Initially, associations between FA and MD of specific tracts were modelled with respect to age, sex and hemisphere using multiple regression. We also included an age*sex interaction term to examine whether age-related trajectories differed for men and women. Linear and quadratic models were compared and, where models with an age^2^ term exhibited a significantly better fit (*p* < 0.05), these results were reported. Due to the large number of comparisons, a threshold of *p* < 0.001 was used to denote significant effects within each model. The age and age^2^ components of these models were illustrated for each tract and water diffusion parameter using kernel density plots to allow clearer visualisation of datapoint overlap than is possible with standard scatter plots with a sample of this size. Tests of the difference between association magnitudes were conducted on the linear component of tract measures, implemented using Williams’ *t* for dependent groups with overlapping correlations (cocor.indep.groups.overlaps) as implemented in the cocor R package (https://cran.r-proiect.org/web/packages/cocor/cocor.pdf).

Confirmatory factor analysis was used to produce one-factor models for each of the five white matter microstructural measurements (FA, MD, ICVF, ISOVF, and OD). On the basis of the first model, for the measurement of FA, we excluded four tracts that had low (<0.3) loadings on the general factor: the Medial Lemniscus, the Middle Cerebellar Peduncle, and the Left and Right Parahippocampal Cingulum. For consistency, we also removed these tracts from the subsequent factor models. In order to improve the model fit for the subsequent use of the factors and the subsequent models, we used the modification indices option in the Mplus v7.3 package^63^ to add additional, residual covariances (paths linking specific tracts to one another). Importantly, none of the results of the study was substantially altered by dropping these residual covariances, or by including the four tracts that were removed. All the models adjusted each tract for sex, age, and—for all measurements other than FA, for which the tracts did not show many appreciable quadratic age curves—age^2^. Fit statistics for the factor models are presented in Supplementary Table 5, standardised factor loadings are provided in Supplementary Table 6, and a list of the residual covariances can be found in Table S7. Factor score estimates were extracted from these models and used in further analyses, and the one-factor model provided the basis for the age-moderation models described below.

We also used structural equation modelling to test whether the age variation in the tracts was best represented as a ‘common pathway’ model (where age has associations with only the latent factor), an ‘independent pathways’ model (where age has associations with each of the individual tracts separately, and not with the latent factor) or a ‘common + independent pathways’ model (where age is associated with the latent factor and also with some specific factors, as required)^64^. To estimate the ‘common + independent pathways’ model, we first included a path from age (which had been centred before inclusion in the model) to the latent factor, then added those paths that produced statistically significant improvements in model fit. For MD, ICVF, ISOVF, and OD, we also included paths from age^2^ alongside every significant age path. We always considered the paths together; if the age path was added, so was the age^2^ path. We then used model fit indices (*χ*^2^ test, Akaike Information Criterion and Bayesian Information Criterion) to examine the fit differences between the three models. Fit statistics are shown in Table S9. In all cases, the ‘common pathways’ model was the worst-fitting. For MD, ICVF, ISOVF, and OD, the ‘common + independent pathways’ model fit either significantly better or no different to the ‘independent pathways model’; for FA, there was some ambiguity, with BIC demonstrating better fit for the ‘common + independent pathways’ model but AIC demonstrating poorer fit.

We examined the degree to which the effect of increasing age on generally lower water diffusion coherence (FA) across tracts was attributable to lower neurite density (ICVF) and/or the amount of tract complexity/fanning (OD) using a multiple mediator model in a structural equation model (SEM) framework^65^ in the lavaan R package^66^. Age was set as the X (independent) variable, the factor score for FA (gFA) was set as the Y (outcome) variable, with gICVF and gOD as two covarying mediators (M). The degree to which the association between X and Y (known as the c path) is attenuated by M (the c’ path) denotes the mediation effect^67^,^68^.

Tests of the relative explanatory power of the factor scores of all five water diffusion parameters to account for age variance – beyond measures more conventionally associated with ageing (total brain, grey matter, white matter and bilateral hippocampal volumes) – was achieved using penalised (elastic net) regression^69^ bootstrapped 1000 times. This method allowed us to identify an optimal combination of predictors from among a group of highly collinear variables. We randomly split the participants into two equal halves (Train and Test). The optimal predictor set was identified by running penalised elastic net regression in the Train group, selecting only those measures that were identified in all 1000 models. A multiple linear regression was then conducted in the Train set and a confirmatory multiple linear regression in the Test set. In general, if the model *β* and *R*^2^ are comparable, the combination of identified predictors are considered optimal for the whole sample.

Age moderation models were estimated separately for each of the white matter microstructural measurements (that is, five models, one for each of FA, MD, ICVF, ISOVF, and OD). The age-moderation models included all of the residual covariances that were added to the one-factor models as described above. In the first set of models, we extended each of the five general factor models to include an interaction parameter representing age moderation of the shared variance across the 22 tracts and an interaction parameter representing age moderation of the tract-specific unique variance components. In other words, we estimated two interaction parameters for each of the 5 diffusion measures. This model can be written for diffusion measure *Y* in tract *t* as:

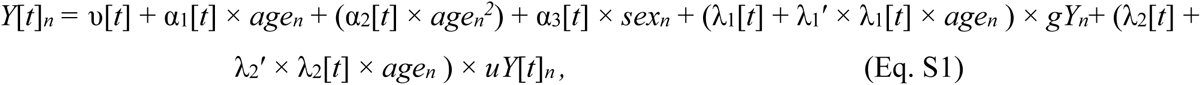

where *v*[*t*] is a tract-specific regression intercept; α_1_[*t*], α_2_[*t*], and α_3_[*t*] are tract-specific regression coefficients for the effects of age, age^2^ (not included for FA), and sex; λ_1_[t] is a tract-specific loading (main effect) on the general factor (*gY*); λ_2_[*t*] is a tract-specific loading (main effect) on the tract-specific unique factor (*uY*[t]); and λ_1_′ and λ_2_′ are tract-invariant interaction parameters representing moderation of the loadings on the general factor and the tract-specific unique factors, respectively. The subscript *n* indicates that a variable varies across individuals. In the above equation, the interaction terms are multiplied by the tract-specific loading in addition to the corresponding latent factor in order to specify age moderation to occur proportionally to the magnitude of the tract-specific loadings (see Cheung *et al*.^70^).

We calculated communality values (proportion of variance explained by the general factor relative to total variance [variance explained by the general factor plus residual variance]^71^, Appendix B,) and their standard errors for 5-year increments of the sample’s age range (that is, for ages 45, 50, 55, 60, 65, 70, and 75 years). Next, using cubic polynomial interpolation, we calculated the expected values at all other ages between 45 and 75 along with their standard errors, and converted these into the age trajectories with 95% confidence intervals shown in Figure 5 in the main document.

To evaluate whether the overall pattern of increasing communality with age was driven by smaller subset of the tracts, we estimated more complex models that estimated tract-specific interaction parameters representing age moderation of loadings on the general factor, and tract-specific interaction parameters representing age moderation of tract-specific uniquenesses^72^. In other words, we estimated 44 interaction parameters (1 interaction for the loading on the general factor and 1 interaction for the unique component, for each of the 22 tracts) for each of the five diffusion measures (FA, MD, ICVF, ISOVF, and OD). This model can be written as

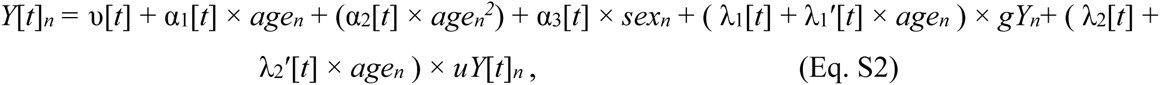

where the interaction terms λ_1_′ and λ_2_′ are estimated individually for each tract, as indicated by the suffix [t]. As interaction terms are estimated individually for each tract, the proportionality constraint (achieved in Eq. S1 via multiplication by the tract-specific loading) is not necessary.

Within the de-differentiation models, we tested whether the age moderation for each loading and each uniqueness was statistically significantly different from zero (that is, whether the parameters λ_1_′ and λ_2_′ differed significantly with age). The results of these tests are provided in Tables S14 and S15 for loadings and uniquenesses respectively. To calculate the communality for each tract (that is, the proportion of the total variance in that tract explained by the factor), we divided the age-specific shared variance in that tract by the age-specific shared-plus-unique (that is, total) variance in that tract^71^ (Appendix B), resulting in the age trend for the communalities shown in the right panels of Figures S8-S12.

Finally, Local Structural Equation Modeling (LOSEM)^73, 74^ was used to provide a non-parametric confirmation of the multi-parameter models described above. Using batch running of Mplus models^75^, we ran 300 one-factor models for each diffusion measure, each aimed at a different part of the age range (300 equal increments between 45 and 75 years). From the outputs of these models, we plotted the factor loadings and uniquenesses as a function of age, for each tract. Again, we also calculated the communality for each tract. The graphical outputs from the LOSEM models are shown in Figures S13 to S17: they confirm the results from the multi-parameter models, providing a more detailed view of the precise age trajectories.

## Acknowledgements

We thank the UK Biobank participants for their participation, and the UK Biobank team for their work in collecting and providing these data for analysis. This research was conducted, using the UK Biobank Resource under approved project 10279, in The University of Edinburgh Centre for Cognitive Ageing and Cognitive Epidemiology (CCACE) (http://www.ccace.ed.ac.uk), part of the cross-council Lifelong Health and Wellbeing Initiative (MR/K026992/1). Funding from the Biotechnology and Biological Sciences Research Council (BBSRC) and Medical Research Council (MRC) is gratefully acknowledged. S.R.C. was supported by MRC grant MR/M013111/1. I.J.D, S.J.R., D.C.L., S.P.H., and G.D. are supported by the Medical Research Council award to CCACE (MR/K026992/1). I.J.D. is additionally by the Dementias Platform UK (MR/L015382/1), and he and S.R.C. by the Age UK-funded Disconnected Mind project (http://www.disconnectedmind.ed.ac.uk). E.M.T.-D. was supported by National Institutes of Health (NIH) research grants HD083613, HD081437, and AA023322. E.M.T.-D. is a member of the Population Research Center at the University of Texas at Austin, which is supported by NIH center grant HD042849. E.M.T.-D. was also supported as a Visiting Scholar at the Russell Sage Foundation. J.M.W. was supported by the Scottish Imaging Network: A Platform for Scientific Excellence (SINAPSE) collaboration (http://www.sinapse.ac.uk).

## Author Contributions

S.R.C. and S.J.R. drafted the manuscript and conducted statistical analyses. E.M.T.-D., M.E.B and I.J.D. supported statistical analysis and edited the manuscript. D.C.L. and S.P.H. provided data support. S.P.H., G.D., J.M.W. and C.R.G edited the manuscript.

